# Synergistic effect of deoxynucleosides and AAV gene therapy for thymidine kinase 2 deficiency

**DOI:** 10.1101/2020.10.08.330969

**Authors:** Carlos Lopez-Gomez, Maria J. Sanchez-Quintero, Eung Jeon Lee, Gulio Kleiner, Jun Xie, Hasan Orhan Akman, Guangping Gao, Michio Hirano

## Abstract

Autosomal recessive thymidine kinase 2 (*TK2*) mutations causes TK2 deficiency, which typically manifests as a progressive and fatal mitochondrial myopathy in infants and children. Treatment with deoxycytidine and thymidine ameliorates mitochondrial defects and extends lifespan of *Tk2* knock-in mouse (TK2^−/−^); however, efficacy is limited by age- and tissue-dependent expression of the cytosolic enzymes Tk1 and Dck. Thus, therapies aimed at systemic restoration of TK2 activity are needed. Here, we demonstrate that delivery of human TK2 cDNA to Tk2^−/−^ mice using AAV9 efficiently rescued Tk2 activity in all the tissues tested except kidney, delayed disease onset, and increased lifespan. Sequential treatment of Tk2^−/−^ mice with AAV9 first followed by AAV2 at different ages allowed us to reduce the viral dose while further prolonging the lifespan. Furthermore, addition of deoxycytidine and deoxythymidine supplementation to AAV9 + AAV2 treated Tk2^−/−^ mice dramatically improved mtDNA copy numbers in liver and kidney, animal growth, and lifespan. These data indicate that combined pharmacological and gene therapies may be highly efficacious for human TK2 deficiency.

## Introduction

Thymidine kinase 2 (TK2), a ubiquitously expressed enzyme critical for the salvage pathways for pyrimidine within mitochondria, phosphorylates both deoxycytidine (dC) and deoxythymidine (dT) to generate deoxycytidine monophosphate (dCMP) and deoxythymidine monophosphate (dTMP). Those deoxynucleoside monophosphates are subsequently phosphorylated to dCTP and dTTP, which are vital building blocks for replication and maintenance of mitochondrial DNA (mtDNA) particularly in post-mitotic cells. Autosomal recessive mutations in the nuclear gene *TK2* cause TK2 deficiency, a rare and devastating form of mtDNA depletion syndrome (MDS). TK2 deficiency manifests most frequently as a progressive and fatal mitochondrial myopathy in infants and children, although around 20% of patients develop the disease during adolescence or adulthood (Garone *et al*, 2018). Studies of *Tk2* homozygous p.His126Arg knock-in (*Tk2*^−/−^) mice indicate that onset and tissue-specificity of the disease are modulated by expression of thymidine kinase 1 (TK1), which catalyze the first step of the cytosolic thymidine salvage pathway (Akman *et al*, 2008, Blazquez-Bermejo *et al*, 2019, Dorado *et al*, 2011).

Treatment of *Tk2*^−/−^ mice with the TK2 substrates, dC and dT, delays disease onset and prolongs the lifespan of the animals by up to 3-fold (Lopez-Gomez *et al*, 2017). Based upon this preclinical study, TK2 deficient patients have been treated with dC+dT in an compassionate use program and have shown improvements in limb weakness, respiratory and swallowing functions, as well as prolonged survival (Dominguez-Gonzalez *et al*, 2019)..

Nevertheless, response to dC+dT therapy in the *Tk2*^−/−^ mice is limited as it only delays onset and slow progression of the disease rather than halting or reversing the course. Several factors constrain the therapeutic response including: 1) rapid degradation of exogenously administered pyrimidine nucleosides by cytidine deaminase and thymidine phosphorylase (TP); 2) restricted delivery of dC and dT via nucleoside transporters into target tissues; and 3) cell-cycle and tissue-specific activities of deoxycytidine kinase (Dck) and thymidine kinase 1 (Tk1) (Blazquez-Bermejo et al, 2019, Lopez-Gomez *et al*, 2019). The nucleoside kinases are particularly critical in *Tk2*^−/−^ mice that develop early central nervous system manifestations, because expression of Tk1 in murine brain is low (Dorado et al, 2011). Because all of these factors limit therapeutic response to dC+dT treatment, treatments aimed at restoring mitochondrial TK2 activity are needed.

Adeno-associated virus (AAV) is being widely used for *in vivo* gene therapy due to its safety profile, transduction efficiency, episomal persistence, and variety of serotypes with different tissue tropism (Wang *et al*, 2019). In fact, clinical trials using AAV vectors have reported a good safety profile, with minimal side-effects, although high doses of AAV has caused liver toxicity (Cukras *et al*, 2018, D’Avola *et al*, 2016, Mendell *et al*, 2017, Mendell *et al*, 2015, Rakoczy *et al*, 2019).

In this study, we demonstrate that transfer of the human TK2 cDNA using AAV9 efficiently rescues TK2 activity in all the tissues assessed except kidney, delayed disease onset, and increased lifespan of *Tk2*^−/−^ mice. Furthermore, sequential treatment with AAV9 first followed by AAV2 at a later age allowed us to reduce the overall viral dose while further prolonging the lifespan of the animals. Additional supplementation of *Tk2*^−/− *AAV9* +*AAV2*^ mice with oral dC+dT further enhanced lifespan, growth, and mtDNA copy numbers in liver and kidney. These finding indicate that the combination of pharmacological and gene therapies is more potent than either therapy alone.

## Results

### Neonatal AAV9 delivery of human TK2 cDNA prolongs lifespan and growth of *Tk2*^−/−^ mice in a dose-dependent manner

Treatment with 4.2×10^10^ vc (low-dose, 2.1-4.2×10^13^ vc/kg) of AAV9-*hTK2* at postnatal day 1 enabled *Tk2*^−/−^ mice to grow normally until day 20 (Fig 1A); however, the animals subsequently developed weakness requiring euthanasia. This treatment significantly prolonged the median lifespan of the *Tk2*^−/−^ mice to 39 days (maximum 52 days) compared to untreated mice with a median survival of 16 days (p=0.0005) This lifespan extension was similar to that observed with high-dose oral nucleoside treatment (dC+dT each at 520 mg/kg/day; *Tk2*^−/−dCdT^) (Lopez-Gomez et al, 2017) (Fig 1B). Unlike *Tk2*^−/−dCdT^ mice, AAV9-*hTK2* treated animals did not develop head tremor.

**Figure 1.**
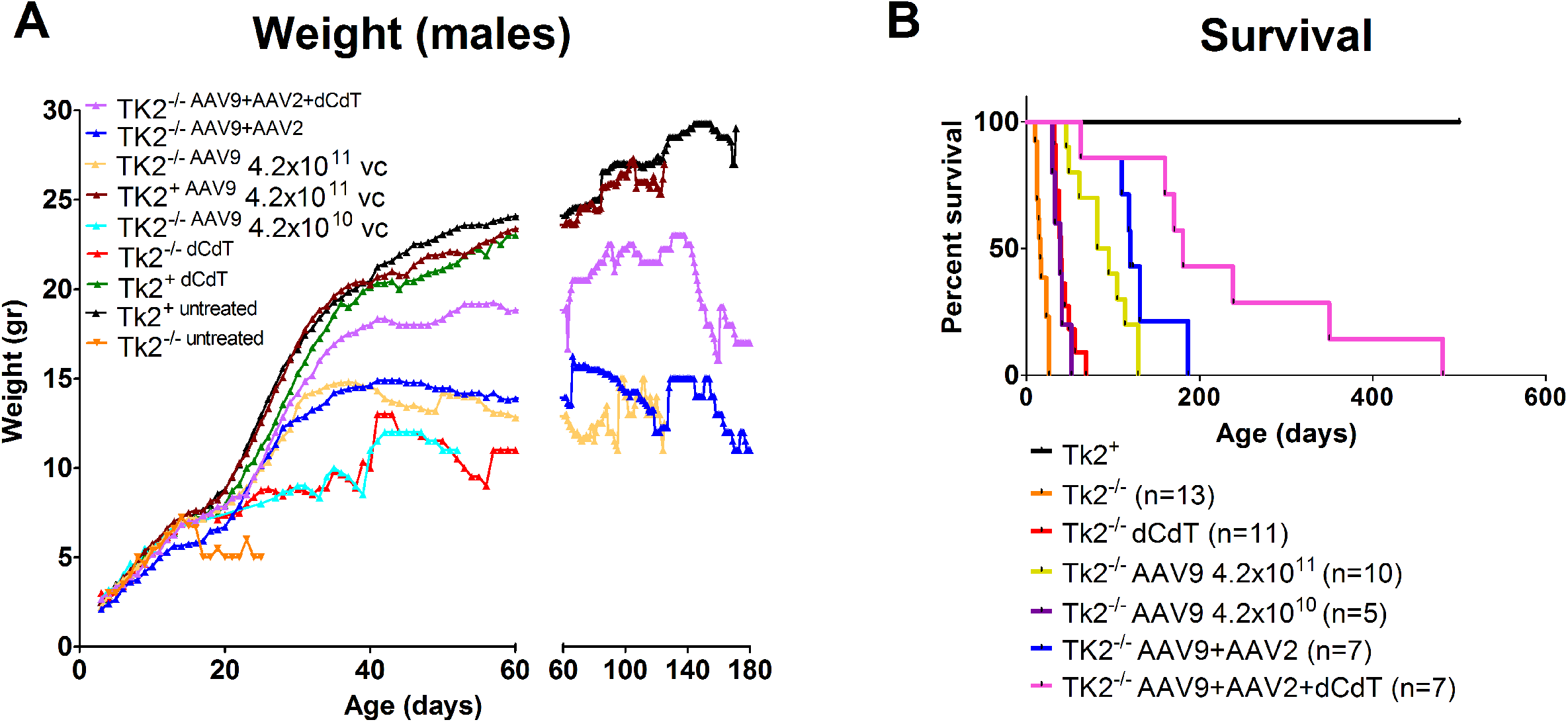
Weight and survival of Tk2^−/−^ with different treatments. Panel A represents average daily weights of male mice for each treatment group. Panel B represents survival as percent within each treatment group.

Increasing the AAV9-*hTK2* dose to 4.2×10^11^ vc (high-dose, 2.1-4.2×10^14^ vc/kg) at postnatal day 1 allowed the mice to grow normally until postnatal day 30 and further extended the median lifespan to 88.5 days (maximum 129 days) relative to untreated *Tk2*^−/−^ mice (p<0.0001) as well as *Tk2*^−/−^ mice treated with low dose (p=0.0003). We found no differences between *Tk2*^+^ and treated *Tk2*^−/−^ mice at age 2 months in terms of motor function, as measured by rotarod test and strength, as measured by grip tests (both bar and grid grip tests) (Fig 2). We did not observe head tremor in the treated mice.

**Figure 2.**
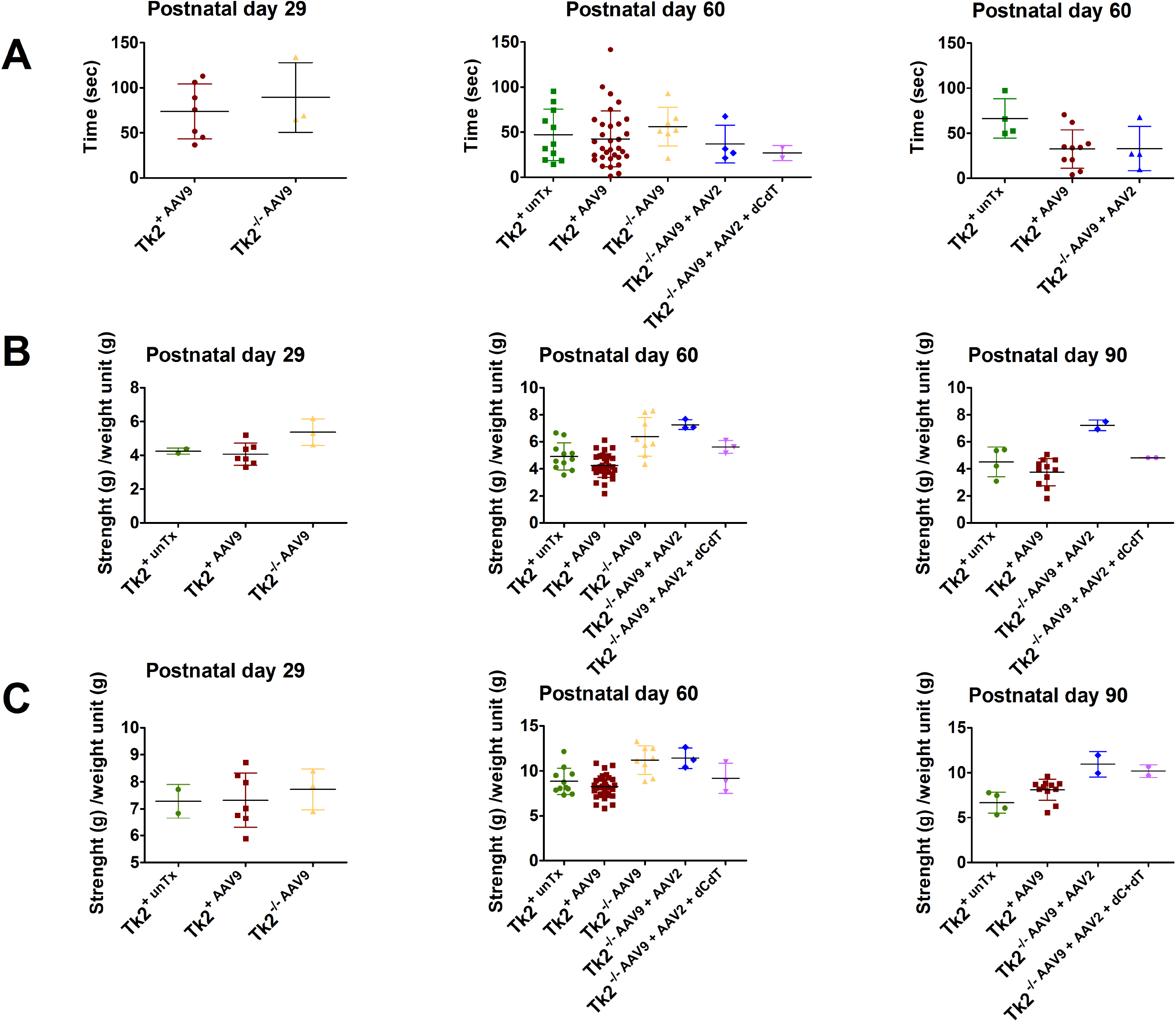
Rotarod and grip tests. Panel A shows results from Rotarod test in male mice. Data is expressed as time to fall from the rotating rod. Each dot represents the average of three test of a single mouse. For each treatment group, average and standard deviation is represented. Panel B shows bar test (upper limbs) and panel C displays grid test (four limbs) in male mice. Data are expressed as strength (measured in g) per weight unit (g). Each symbol represents the average of three tests of a single mouse. For each treatment group, average and standard deviation are represented.

### AAV9-*hTK2* rescues Tk2 activity and mtDNA depletion in muscle, brain, and liver, but not kidney

Mice treated with high-dose AAV9-*hTK2* at postnatal day 1 (*Tk2*^+*AAV9*^ and *Tk2*^−/− *AAV9*^ combined) showed widespread *TK2* mRNA expression, which was sustained up through age 18 months (Table 1). The highest *TK2* transcript levels were observed in skeletal muscle (>10^4^-fold above endogenous murine *Tk2*) followed by brain (>10^3^-fold) and liver (>6-fold). In contrast, expression of human TK2 in kidney was less robust, showing levels of 265% ± 145% at age 1 month, declining to 34% ± 26% at 2 months and 18% ± 6% at 6 months.

**Table 1.** mRNA expression of human TK2.

TK2 mRNA expression was calculated as percentage of expression of endogenous mouse *Tk2*.
| Tissue | 1 month (n=7) | 2 months (n=15) | 6 months (n=6) | 18 months (n=11) |
| --- | --- | --- | --- | --- |
| Brain | $2.1 \times 10^3 \pm 1.8 \times 10^3$ | $3.7 \times 10^3 \pm 3.0 \times 10^3$ | $2.1 \times 10^3 \pm 2.9 \times 10^3$ | $1.8 \times 10^3 \pm 2.8 \times 10^3$ |
| Liver | $864.7 \pm 271.0$ | $1.2 \times 10^3 \pm 327.3$ | $825.3 \pm 701.4$ | $636.2 \pm 243.0$ |
| Kidney | $265.1 \pm 145.4$ | $34.3 \pm 25.7$ | $17.7 \pm 6.2$ | |
| Muscle | $6.9 \times 10^3 \pm 6.6 \times 10^3$ | $3.6 \times 10^4 \pm 1.7 \times 10^4$ | $3.7 \times 10^4 \pm 1.9 \times 10^4$ | $1.6 \times 10^5 \pm 7.3 \times 10^4$ |

Concordant with the increases in *TK2* transcript, mean TK2 activity from *Tk2*^−/− *AAV9*^ mice was rescued at postnatal day 29 in muscle (40-fold elevated TK2 activity relative to Tk2 level in *Tk2*^+^ mice), brain (130%), and liver (93%), but not in kidney (37%) (Fig 3A and supplemental Table 1). At P60, TK2 activity in muscle of *Tk2*^−/− *AAV9*^ mice remained high (33-fold elevated), but decreased to 53% in brain and was stable in liver (84%) and kidney (41%) (Fig 3B). These results demonstrate efficiency of the designed AAV9-*hTK2* vector to transduce a human protein in the two main tissues affected in our mouse model, brain and muscle, and confirm the poor transduction efficiency of the vector in kidney.

**Figure 3.**
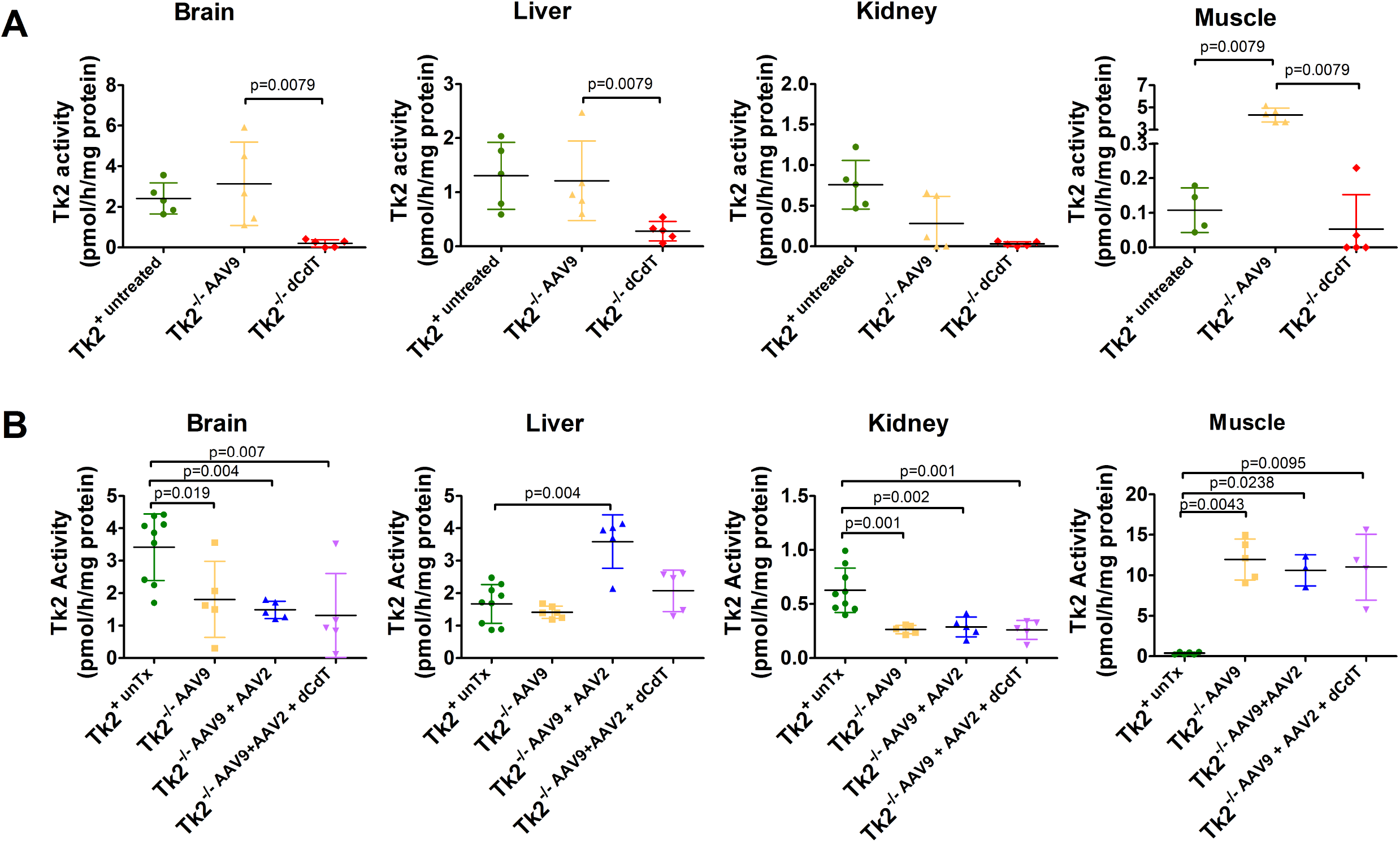
TK2 activity. TK2 activity from mice at age 29 (panel A) and 60 days (panel B). Data are expressed as pmol of product generated per hour and normalized to mg of protein for each sample. Each symbol represents the value obtained for each tissue in individual mice. For each treatment group, average and standard deviation are represented. P values are the results of Mann-Whitney tests.

Expression of TK2 in P29 *Tk2*^−/− *AAV9*^ mice significantly increased levels of mtDNA relative to *Tk2*^−/− *dCdT*^ in brain, heart, liver, and muscle (Fig 4A). In kidney, mtDNA levels were mildly increased in *Tk2*^−/− *AAV9*^ (47.4±7.7%) compared to animals treated with dC+dT (32.8 ± 3.8%) [p=0.016]) but significantly lower than in wild-type mice (100.0 ± 10.3% [p=0.008]). At age P60 (2 months), mtDNA levels in *Tk2*^−/− *AAV9*^ were moderately reduced in brain (80.8%), heart (77.3%), muscle (70.2%) and liver (67.3%) but severely depleted in kidney (12.7%) (Fig 4B).

**Figure 4.**
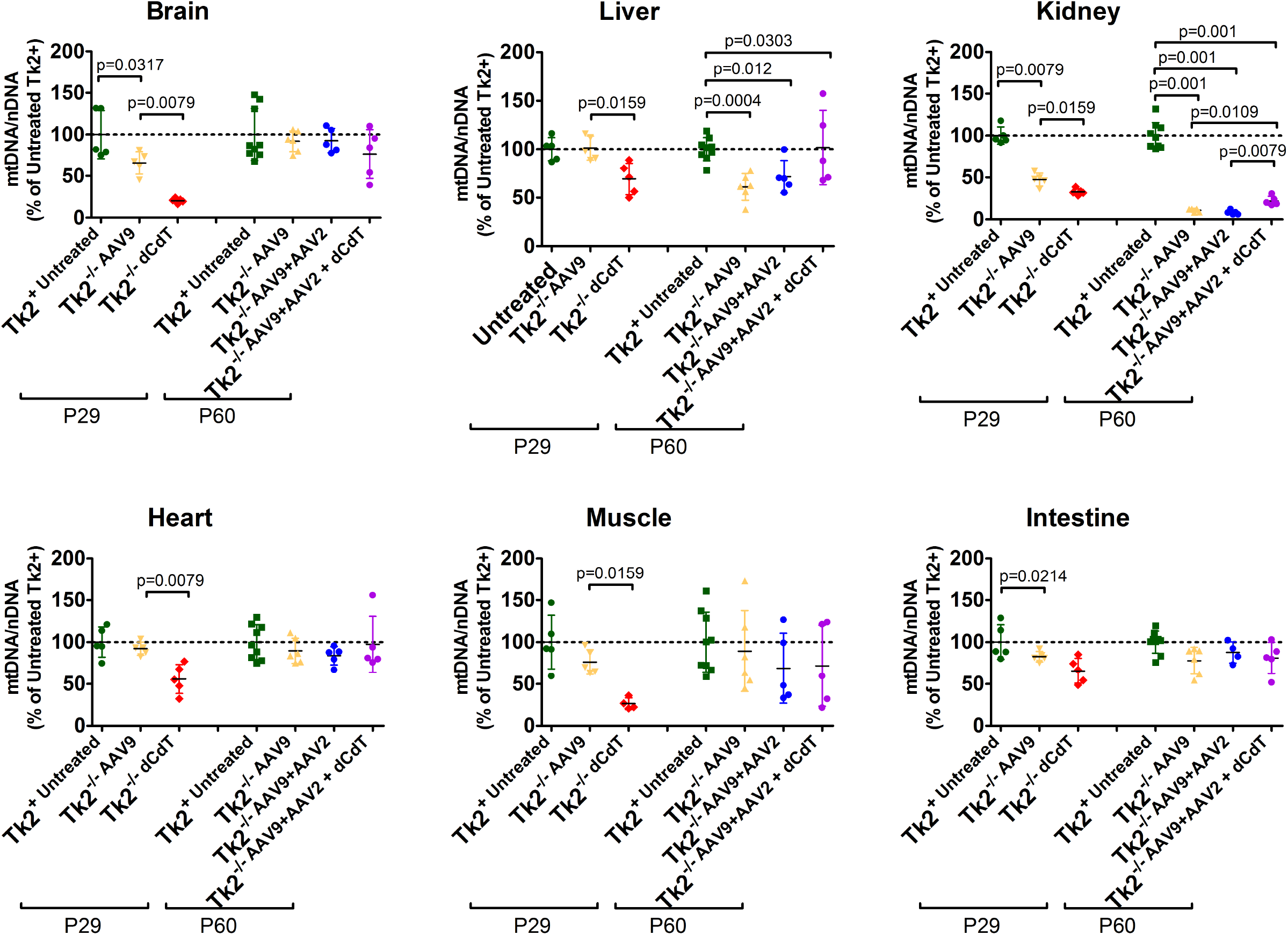
mtDNA levels. Data are expressed as percents of mtDNA in untreated Tk2^+^ mice. Each symbol represents the average of three measurements in a single tissue of individual mice. For each treatment group, average and standard deviation are represented. P values are the results of Mann-Whitney tests.

### mtDNA depletion in kidney is associated with renal dysfunction

At age P60, *Tk2*^−/− *AAV9*^ mice showed higher levels of blood urea nitrogen (BUN) compared to *Tk2*^+ *AAV9*^ mice (68.7 ± 18.2 mg/dL *vs* 30.8 ± 8.9 mg/dL [p=0.009]) (Fig 5 and Tables 2 and 3). Furthermore, in 2 out of 6 *Tk2*^−/− *AAV9*^ mice, high levels of BUN were accompanied by elevated plasma creatinine (>0.3 mg/dL; normal <0.2 mg/dL) indicating impaired renal function. Moreover, one *Tk2*^−/− *AAV9*^ mouse at end-stage (age P129) showed plasma levels of BUN (>140 mg/dL) and creatinine (>0.5 mg/dL) above detection limits indicating that kidney dysfunction was progressive and contributing to the early mortality of the *Tk2*^−/− *AAV9*^ mice.

**Figure 5.**
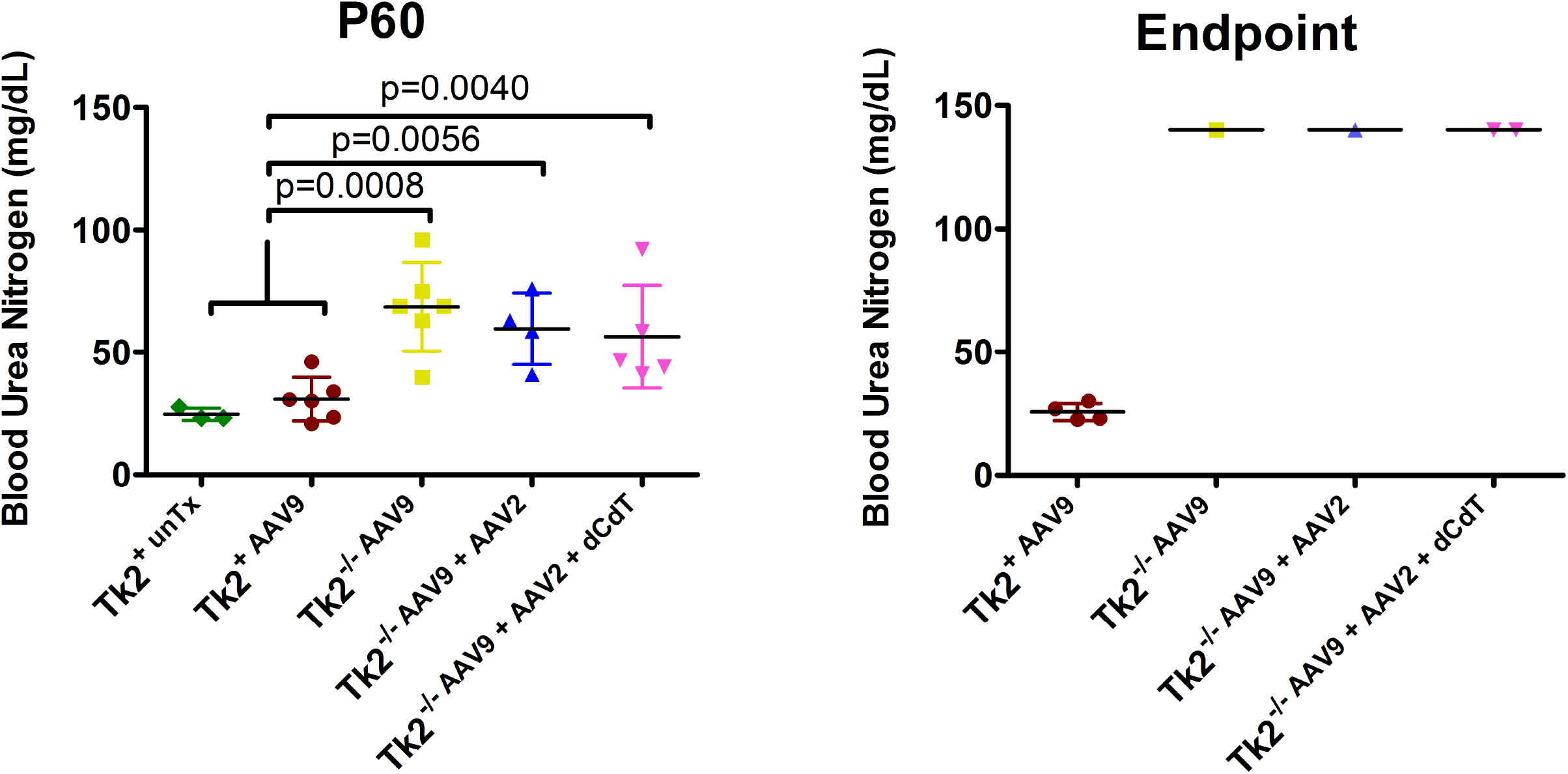
Blood Urea Nitrogen levels. Data is expressed as mg/dL of plasma. Each symbol represents the value for a single mouse. Left panel represents BUN at age 60 days. For each treatment group, average and standard deviation are represented. P values are the results of Mann-Whitney tests. Right panel represents BUN at endpoint. Values from all the Tk2^−/−^ mice at endpoint were in fact above the detection limit of the technique (>140mg/dL).

**Table 2.** Creatinine levels in plasma at age 60 days.

Values indicate total number of mice from each treatment group showing a determined level of creatinine.
| Creatinine | Tk2<sup>+</sup> untreated | Tk2<sup>+</sup> AAV9 | Tk2<sup>-/-</sup> AAV9 | Tk2<sup>-/-</sup> AAV9 + AAV2 | Tk2<sup>-/-</sup> AAV9+AAV2 + dCdT |
| --- | --- | --- | --- | --- | --- |
| <0.2 mg/dL | 2 | 0 | 0 | 2 | 0 |
| 0.2 mg/dL | 0 | 5 | 4 | 1 | 0 |
| 0.3 mg/dL | 1 | 1 | 0 | 1 | 3 |
| >0.3 mg/dL | 0 | 0 | 2 | 0 | 2 |
| Total mice assessed | 3 | 6 | 6 | 4 | 5 |

**Table 3.** Creatinine levels in plasma at end-stage.

Values indicate total number of mice from each treatment group showing a determined level of creatinine.
| Creatinine | Tk2<sup>+</sup> AAV9 | Tk2<sup>-/-</sup> AAV9 | Tk2<sup>-/-</sup> AAV9 + AAV2 | Tk2<sup>-/-</sup> AAV9+AAV2 + dCdT |
| --- | --- | --- | --- | --- |
| 0.2 mg/dL | 4 | 1 | 1 | 1 |
| 0.3 mg/dL | 0 | 0 | 0 | 0 |
| >0.3 mg/dL | 0 | 1 | 0 | 1 |
| Total mice assessed | 4 | 2 | 1 | 2 |

### Treatment with AAV9-hTK2 followed by redosing with AAV2-hTK2 prolongs survival of *Tk2*^−/−^ mice and partially ameliorates renal functions

To enhance expression of human *TK2* gene in kidney and further improve AAV9-*hTK2* therapy, we assessed the effects of subsequent treatment with AAV2-*hTK2*. A single dose of 2.1×10^11^ vc (1.05-2.1×10^14^ vc/kg) of AAV9-*hTK2* was administered at P1, followed by a second treatment with 1.05×10^11^ vc (0.7-1.8×10^13^ vc/kg) of AAV2-*hTK2* administered at P29. This treatment regime results in a lower total viral dose (3.15×10^11^ vc; 1.12-2.3×10^14^ vc/kg) relative to high single neonatal dose regime of AAV9-*hTK2* (4.2×10^11^ vc; 2.1-4.2×10^14^ vc/kg).

*Tk2*^−/− *AAV9*+*AAV2*^ mice showed significantly longer survival compared to high-dose *Tk2*^−/− *AAV9*^ mice (median survival of 120 days, maximum 187 days, versus 88.5 days [p=0.045]) (Fig 1) although growth curves are very similar in both treatment groups (Fig 2). To assess whether this longer survival is the consequence of improved renal function, we assessed renal glomerular function by measuring protein levels in urine (Table 4). Between ages 21 and 29 days, 2 out of 4 *Tk2*^−/− *AAV9*^ mice showed protein levels >500 mg/dL, indicating glomerulopathy. A third mouse had 30 mg/dL protein in urine, while a fourth mouse showed only trace levels of protein, similar to *Tk2*^+^ mice. The three mice with increased proteinuria were treated with AAV2-*hTK2* at age 29 days (*Tk2*^−/− *AAV9*+*AAV2*^ mice), while the fourth was not treated further. At age 60 days, the three *Tk2*^−/− *AAV9*+*AAV2*^ mice had only trace urine protein, while the fourth *Tk2*^−/− *AAV9*^ mouse showed levels > 500 mg/dL. With improvements in proteinuria, BUN levels in *Tk2*^−/− *AAV9*+*AAV2*^ mice were mildly reduced compared to *Tk2*^−/− *AAV9*^ mice (59.7±14.6 *vs* 68.7±18.2 mg/dL, p>0.05), but significantly higher than in *Tk2*^+^ mice (30.8±8.9 mg/dL, p=0.006). At age 60 days, serum creatinine was also elevated (>0.3mg/dL) in 2 of 6 *Tk2*^−/− *AAV9*^ mice, but not in the 4 *Tk2*^−/− *AAV9*+*AAV2*^ or 3 *Tk2*^+^ mice. In most tissues, TK2 activities in *Tk2*^−/− *AAV9*+*AAV2*^ mice at age 60 days were similar to Tk2 activities in *Tk2*^−/− *AAV9*^ mice (Fig 3B) confirming that transduction of hTK2 to kidney had not improved significantly. Furthermore, mtDNA copy number were similar in both *Tk2*^−/− *AAV9*^ and *Tk2*^−/− *AAV9*+*AAV2*^ mice in all tissues, except kidney, which showed severe mtDNA depletion in *Tk2*^−/− *AAV9*+*AAV2*^ mice at age 60 days (kidney mtDNA level 8.2 ± 2.6% vs 100.0 ± 15.8% in *Tk2*^−/− *AAV9*+*AAV2*^ vs Tk2^+^ mice [p=0.001] and 10.7 ± 2.2% in *Tk2*^−/− *AAV9*^ mice [not significant] (Fig 4). In aggregate, correction of proteinuria after AAV2 treatment may be indicative of improved TK2 transduction in kidney glomeruli (due to either different tropism of AAV2 vs AAV9 or age of treatment), but low mtDNA levels in whole kidney suggests that TK2 transduction remains low in most of kidney cell types and therefore insufficient to fully rescue renal functions.

**Table 4.** Protein levels in urine.

| Mouse genotype | Treatment before P21-P29 | Treatment after P29 | Protein level |  |
| --- | --- | --- | --- | --- |
|  |  |  | P21-P29 | P60 |
| Tk2+ #1 | None | None | Trace | Trace |
| Tk2-/- #1 | AAV9-hTK2 | AAV2-hTK2 | 500 mg/dL | Trace |
| Tk2-/- #2 | AAV9-hTK2 | AAV2-hTK2 | 500 mg/dL | Trace |
| Tk2+ #2 | None | None | Trace | Trace |
| Tk2+ #3 | None | None | Trace | Trace |
| Tk2-/- #3 | AAV9-hTK2 | AAV2-hTK2 | 30 mg/dL | Trace |
| Tk2+ #4 | None | None | Trace | Trace |
| Tk2-/- #4 | AAV9-hTK2 | None | Trace | 500 mg/dL |

### Supplementation with oral dC+dT further improves effects of gene therapy and prolongs lifespan

A set of *Tk2*^−/− *AAV9*+*AAV2*^ mice was further supplemented with 520 mg/kg/day of oral dC+dT from age P21. Survival of *Tk2*^−/− *AAV9*+*AAV2*+*dCdT*^ mice was significantly prolonged (median survival of 181 days and maximum 481 days) compared to survival with other treatments (*Tk2*^−/− *AAV9*^ [p=0.0013] and *Tk2*^−/− *AAV9*+*AAV2*^ [p=0.062]) (Fig 1).

Growth also improved in *Tk2*^−/− *AAV9*+*AAV2*+*dCdT*^ mice. While all three treatments (*Tk2*^−/− *AAV9*^, *Tk2*^−/− *AAV9*+*AAV2*^ and *Tk2*^−/− *AAV9*+*AAV2*+*dCdT*^ mice) increased weight to ~ 80% of *Tk2*^+^ mice at age 29 days (Fig 10), both *Tk2*^−/− *AAV9*^ and *Tk2*^−/− *AAV9*+*AAV2*^ mice subsequently stopped gaining weight (Fig 2). In contrast, *Tk2*^−/− *AAV9*+*AAV2*+*dCdT*^ mice at age 60 days weighed significantly more (82.3±11.4%) than *Tk2*^−/−^ with any other treatments (55.7±11.3% in Tk2^−/− *AAV9*^ mice [p=0.0001] and 57.4±10.5% in *Tk2*^−/− *AAV9*+*AAV2*^ mice [p=0.0002]) (Fig 1).

As expected, TK2 activity at age 60 days was similar in both *Tk2*^−/− *AAV9*+*AAV2*+*dCdT*^ and *Tk2*^−/− *AAV9*+*AAV*2^ mice in all the tissues (Fig 3B) with the exception of liver, which was lower in mice supplemented with deoxynucleosides (2.08 ± 0.64 pmol/h/mg protein in *Tk2*^−/− *AAV9*+*AAV2*+*dCdT*^ mice *vs* 3.59 ± 0.82 in *Tk2*^−/− *AAV9*+*AAV2*^ mice, p=0.056). Levels of mtDNA were similar in tissues from *Tk2*^−/− *AAV9*+*AAV2*+*dCdT*^ and *Tk2*^−/− *AAV9*+*AA*V2^ mice, except in liver and kidney. Deoxynucleoside supplementation produced higher levels of mtDNA in liver (101.7 ± 38.3% in *Tk2*^−/− *AAV9*+*AAV2*+*dCdT*^ mice vs 71.6 ± 16.9% in *Tk2*^−/− *AAV9*+*AAV2*^ mice [p>0.05] and 61.0 ± 13.7% in *Tk2*^−/− *AAV9*^ mice [p=0.030] and in kidney (22.1 ± 5.4% in *Tk2*^−/− *AAV9*+*AAV2*+*dCdT*^ mice vs 8.2 ± 2.6% in *Tk2*^−/− *AAV9*+*AAV2*^ mice [0.0079] and 10.7 ± 2.2% in *Tk2*^−/− *AAV9*^ mice [p=0.0303]. The slightly higher levels of mtDNA in kidney partially rescued renal function in *Tk2*^−/− *AAV9*+*AAV2*+*dCdT*^ which showed increased BUN similar to *Tk2*^−/− *AAV9*^ and *Tk2*^−/− *AAV9*+*AAV2*^ mice at age 60 days and at end-stage as well as elevated creatinine (>0.3 mg/dL) in 2 of 5 mice at age 60 days and 1 of 2 mice at end-stage.

## Discussion

Mitochondrial DNA Depletion Syndrome (MDS) encompasses a heterogeneous group of autosomal recessive mitochondrial disorders characterized by severe reduction of mtDNA copy number. The causative gene largely dictates organ involvement with myopathy (*TK2*), encephalomyopathy (*SUCLA2*, *SUCLG1*, or *RRM2B*), encephaloneurogastrointestinal (*TYMP*) or hepatoencephalopathy (*DGUOK*, *MPV17*, *POLG1*, or *C10orf2*) phenotypes (El-Hattab & Scaglia, 2013). To date, there are no cures for MDS and disease-modifying treatments are only available for only the forms caused by pathogenic variants in *TYMP* causing mitochondrial neurogastrointestinal encephalopathy (MNGIE) (Bax *et al*, 2019, Halter *et al*, 2011, Kripps *et al*, 2020, Nishino *et al*, 1999, Yadak *et al*, 2017) and *TK2* leading to TK2 deficiency (Dominguez-Gonzalez et al, 2019).

Treatment of TK2 deficiency with deoxynucleoside has showed promising results in both knock-in and knock-out Tk2 mouse models (Blazquez-Bermejo et al, 2019, Lopez-Gomez et al, 2017), as well as in patients receiving compassionate use therapy (Dominguez-Gonzalez et al, 2019). In the study by Dominguez-Gonzalez *et al* (Dominguez-Gonzalez et al, 2019), the authors observed amelioration of symptoms and prolonged survival of patients with open-label dC+dT therapy compared to historical controls. Based upon these observations, a Phase 2 prospective, open-label clinical trial has been initiated to further assess the safety and efficacy of deoxynucleoside treatment in TK2 deficient patients (NCT03845712). Nevertheless, results in Tk2 deficient mice showed limited efficacy, likely due to several factors tempering response to dC+dT (Blazquez-Bermejo et al, 2019, Lopez-Gomez et al, 2019). Potency of dC+dT therapy may be limited in encephalomyopathic forms of human TK2 deficiency, because low expression of TK1 in brain may contrain response of the encephalopathy to this therapy. Therefore, more effective treatments are needed for TK2 deficiency as well as other forms of MDS.

In this study, we have assessed effects of AAV9 delivery of the human TK2 cDNA to our Tk2^−/−^ mice. We selected AAV9 because of its wide tropism and we incorporated chicken beta actin promoter to enhance its expression in muscle, which is the most affected tissue in patients. Our results showed that efficiency of transduction was very high, especially in skeletal muscle. Furthermore, expression of the human TK2 mRNA was stable up to 18 months after the treatment, with the exception of kidney, which downregulated the expression of hTK2 from more than 200% to less than 50% of endogenous murine Tk2.

Because TK2 is an enzyme rather than a structural protein and because heterozygous *TK2* mutation carriers are asymptomatic, we hypothesized that partial activity should be sufficient to achieve normal levels of mtDNA. In fact, mtDNA levels in both muscle and brain from *Tk2*^−/−*AAV9*^ mice at age 60 days were similar to those observed in Tk2^+^ mice, although brain showed about half of the TK2 activity of the Tk2^+^ mice, which confirms that complete rescue of the TK2 activity is not necessary to achieve a therapeutic response. In contrast, kidney from *Tk2*^−/−*AAV9*^ mice, which also showed about half of the activity of Tk2^+^ mice, showed a devastating mtDNA depletion that led to kidney dysfunction. Differential transduction of the human TK2 gene in different kidney cell types, with few cells acquiring many copies of the transgene while most remaining untransfected, may explain the discrepancy between the moderate TK2 activity in the whole tissue and the low levels of mtDNA observed.

Although treatment with AAV9-*hTK2* did not work as effectively in kidney relative to other tissues, disease onset was delayed, disease course improved, and lifespan was prolonged from a median of 16 days to 88.5 days. In fact, none of the *Tk2*^−/−*AAV9*^ mice showed head tremor at endpoint, which is a typical sign of disease in this mouse model; the absence of tremor indicates amelioration of the encephalopathy.

It has been previously reported that neonatal treatment is not efficient in transducing kidney and liver tissues in mice (Bostick *et al*, 2007), which may explain the poor transduction of hTK2 observed in kidney, although transduction in liver was better than reported. Therefore, to improve this therapy, we modified our treatment plan with a second injection at age 29 days, using a different serotype to avoid immunogenicity. We chose AAV2 serotype because of its reported high transduction efficiency in mouse kidney (Takeda *et al*, 2004) as well as its distinctive serological property to AAV9. Interestingly, using this dual AAV treatment with total vc 75% of the high-dose AAV9 alone, TK2 activity in tissues remained at the same levels as in high-dose *Tk2*^−/−*AAV9*^ mice. TK2 activity in kidney was not detectably improved and renal dysfunction was only partially ameliorated (decreased proteinuria and slightly reduced BUN levels in plasma), lifespan of *Tk2*^−/−*AAV9*+*AAV2*^ mice was prolonged to a median of 119 days (34% increase compared to high-dose *Tk2*^−/−*AAV9*^ mice). *Tk2*^−/−*AAV9*+*AAV2*^ weight and course of disease did not improve relative to high-dose *Tk2*^−/−*AAV9*^ mice and there were no differences in the levels of mtDNA that could explain the increased survival between *Tk2*^−/−*AAV9*^ and *Tk2*^−/− *AAV9*+*AAV2*^ mice. Importantly, the AAV9+AAV2 combination produced the same effect on mtDNA levels and even improved survival using a lower overall viral dose compared to high-dose AAV9 alone. Hence, use of different AAV serotypes at different time points is a promising strategy to enhance this gene therapy for TK2 deficiency.

To further improve this treatment, we supplemented *Tk2*^−/−*AAV9*+*AAV2*^ mice with 520 mg/kg/day of oral dC+dT from day 21 in order to enhance TK2 activity in tissues with low expression of the human TK2 transgene. This dC+dT supplementation further prolonged the median lifespan to 481 days. Interestingly, weight of *Tk2*^−/−*AAV9*+*AAV2*+*dCdT*^ mice was also significantly increased, which may be related to a significantly increased mtDNA copy number in liver and kidney relative to *Tk2*^−/−*AAV9*^ mice. Although TK2 activity in liver was slightly higher in *Tk2*^−/−*AAV9*+*AAV2*^ and *Tk2*^−/−*AAV9*+*AAV2*+*dCdT*^ relative to *Tk2*^−/−*AAV9*^ mice, which confirms that in tissues with TK2 activity, supplementation with dC+dT improves the replication and maintenance of mtDNA.

In conclusion, our study provides the first demonstration of efficacy of AAV-mediated gene therapy for Tk2 deficiency in a mouse model. Systemic transduction of the human *TK2* cDNA using AAV9 is highly effective in targeting skeletal muscle, the most affected tissue in patients with TK2 deficiency. Furthermore, the use of the chicken beta actin promoter in our system proved to be highly efficient in enhancing transgene expression and activity of TK2 in most tissues, but most effectively in skeletal muscle. Finally, we propose dC+dT supplementation as a co-treatment to enhance the effects of TK2 gene therapy and perhaps reduce the viral dose, which may reduce known side-effects of high doses this gene therapy including hepatotoxicity.

## Methods

### Vector construction and production

Thymidine Kinase 2 Human Untagged Clone (NM_004614) containing the cDNA for transcript variant 1 was purchased from OriGene Technologies, Inc. (Rockville MD) (supplemental information 1). The 1016 bp TK2 cDNA containing the full-length coding sequence was extracted by *Eag*I digestion. A pAAVsc CB6 PI vector described by Rashonejad and colleagues (Rashnonejad *et al*, 2016) was used to generate AAV9 virus containing TK2 cDNA. After generating blunt ends by treating vector and insert DNA with T4 DNA polymerase (New England Biolabs), we used blunt-end ligation to insert TK2 cDNA into *Cla*I digested pAAVsc CB6 PI vector (supplement information 2).

### Mice

Generation and characterization of *Tk2 H126N* knock-in mice was previously reported (Akman et al, 2008). All experiments were performed according to a protocol approved by the Institutional Animal Care and Use Committee of the Columbia University Irving Medical Center, and are consistent with the National Institutes of Health Guide for the Care and Use of Laboratory Animals. Mice were housed and bred according to international standard conditions, with a 12-hour (h) light, 12-h dark cycle.

### Phenotype assessment

Body weight was assessed daily, because we previously observed that cessation of weight gain is the initial phenotypic sign of disease followed by head tremor (Akman et al, 2008). To define the degree of safety and efficacy of each therapy, we recorded age-at-onset of disease, types and severity of manifestations, side effects, proportion of treatment termination due to adverse events, and survival duration in treated and untreated *Tk2*^−/−^ mice. Behavior, survival time, and body weights of the mice were assessed daily beginning at postnatal day 2.

Motor function was assessed with an accelerating rotarod performance test (Economex Rota-Rod, Columbus Instruments). Four mice were tested simultaneously on a 3.5-cm rotating rod with each mouse separated by a 3 mm wide × 60 cm high opaque Plexiglas wall. The rotarod was started at 10 rpm and accelerated by 2.5 rpm every 10 seconds (s). After a training phase of three trials, three motor performances for each mouse were averaged and analyzed.

Grip strength was measured using a Grip Strength Meter (Columbus Instruments). Mice were allowed to grip a bar with the upper limbs and a rectangular grid with all four limbs, followed by pulling the mice until they released; three force measurements were recorded in each trial. Strength, measured by mass units (grams [g]), was normalized to whole body weight of each animal (g).

### Treatment administration and experimental plan

Two cohorts of animals composed by both *Tk2*^+^ and *Tk2*^−/−^ mice were treated with either 4.2×10^10^ or 4.2×10^11^ viral copies (vc) of AAV9-*hTK2* in a total volume of 35 μL at postnatal day 1 intravenously (retro-orbital injection).

A third cohort of animals was treated with 2.1×10^11^ vc of AAV9-*hTK2* in a total volume of 35 μL at postnatal day 1 intravenously (retro-orbital injection), and then treated with 1.05×10^11^ vc of AAV2-*hTK2* in a total volume of 100 μL intravenously (tail vein injection). A subgroup of this cohort was supplemented with dC and dT (520mg/kg/day each) (Hongene Biotechnology, Inc.) in the drinking water, assuming a daily water consumption of 4 mL per mouse.

In addition, we treated *Tk2*^+^ and *Tk2*^−/−^ mice with oral dC and dT (520mg/kg/day each) from postnatal day 4 to assess survival and molecular and biochemical assessments in mice treated with only deoxynucleosides at ages 13 and 29 days.

### Human TK2 gene expression

RNA from different tissues was isolated using Trizol™ reagent (Thermo Fisher Scientific) following the manufacturer protocol. cDNA was synthesized from 500 ng of RNA using SuperScript™ VILO™ cDNA Synthesis Kit (Invitrogen™) following manufacturer protocol. Real-time PCR was performed in a Step One Plus Real Time PCR System (Applied Biosystems) using a Taqman probe specific for the human *TK2* transcript (Hs00936914, Thermo Fisher Scientific) and a Taqman probe specific for the murine *Tk2* transcript (Mm01250904, Thermo Fisher Scientific). Expression of murine *GAPDH* (Mm99999915, Thermo Fisher Scientific) was used as endogenous control. Data was analysed using the ddCt method and expression of human *TK2* was calculated as percentage relative to the endogenous mouse *Tk2*.

### TK2 activity

TK2 activity was measured using tritium-labeled bromovinyl deoxyuridine as previously described (Franzolin *et al*, 2006), with the following modifications: DEAE filtermat filters (PerkinElmer, Waltham, MA, USA) were used and washed twice for 5 minutes in 1mM ammonium formate and once for 5 minutes in H_2_O. TK2 enzyme activity is expressed in pmol/hour/mg of protein.

### Mitochondrial DNA quantification

Real-time PCR was performed with the primers and probes for murine COX I gene (mtDNA) and glyceraldehyde-3-phosphate dehydrogenase (Gapdh, nuclear DNA [nDNA]) (Applied Biosystems, Invitrogen, Foster City, CA, USA) as described using ddCt method in a Step One Plus Real Time PCR System (Applied Biosystems) (Dorado et al, 2011). mtDNA values were normalized to nDNA values and expressed as a percentage relative to wild-type (100%).

### Immunoblotting

To assess protein levels of the mitochondrial respiratory chain complexes, thirty micrograms of whole brain extracts were electrophoresed in 10-20% Tris-Glycine Gel (Novex™ WedgeWell™, Invitrogen) and transferred to Immun-Blot™ PVDF membranes (Biorad, Hercules, CA, USA). Membranes were probed with MitoProfile^®^ Total OXPHOS Rodent WB Antibody Cocktail of antibodies (MitoSciences, Eugene, OR, USA) against complex I (CI) subunit Ndufb8, complex II (CII) 30kDa, complex III (CIII) Core protein 2, complex IV (CIV, COX) subunit I, and complex V (CV) alpha subunit.

Levels of Vinculin were assessed as loading control (anti-Vinculin AB18058). Protein– antibody interaction was detected with either anti-mouse or anti-rabbit peroxidase-conjugated mouse IgG antibody (Sigma-Aldrich, St Louis, MO, USA), using Amersham™ ECL Plus western blotting detection system (GE Healthcare Life Sciences, UK). Quantification of proteins was carried out using NIH ImageJ 1.37V software. Average grey value was calculated within selected areas as the sum of the grey values of all the pixels in the selection divided by the number of pixels.

### Mitochondrial respiratory chain enzyme activities by spectrophotometer analysis

Mitochondrial RC enzymes analysis was performed in cerebrum tissue as described (Birch-Machin *et al*, 1994, DiMauro *et al*, 1987).

### Kidney function

Kidney function was assessed measuring blood urea nitrogen (BUN) levels in plasma in a Heska Element DC analyzer, following manufacturer procedures. Additionally, we assessed levels of protein in mice urine using Chemstrip^®^ 10 (Roche Diagnostics GmgH, Mannheim, Germany).

### Statistical Analysis

Data are expressed as the mean ± SD of at least 3 experiments per group. For data grouped in columns, Mann-Whitney test was used to compare each group. For survival curves, data is expressed as median ± SD and Mantel-Cox test was used to compare groups. A *p*-value of <0.05 was considered to be statistically significant.

## Data Availability

All primary data are available upon request.

## Acknowledgements

The authors would like to acknowledge Professor Liya Wang for assistance with the assessment of TK2 activity, and Maoxue Tang, PhD, for assistance with the retro-orbital injection to mice.

This work was supported by a research grant from the NIH (P01 HD32062) (MH). MH is supported by the Arturo Estopinan TK2 Research Fund, Nicholas Nunno Foundation, JDM Fund for Mitochondrial Research, Shuman Mitochondrial Disease Fund, and the Marriott Mitochondrial Disease Clinic Research Fund (MMDCRF) from the J. Willard and Alice S. Marriott Foundation. MH also acknowledges support from NIH U54 NS078059 from NINDS and NICHD.

## Conflicts of Interest

Columbia University has a patent for deoxynucleoside therapies for mitochondrial DNA depletion syndrome including TK2 deficiency, which is licensed to Modis Therapeutics a wholly owned subsidiary of Zogenix Inc.; this relationship is monitored by an unconflicted external academic researcher. Dr. Hirano is a co-inventor of this patent. CUIMC has received royalty payments related to the development and commercialization of the technology; Dr. Hirano has received shares of the royalty payments following Columbia University policies. MH is a paid consultant to Modis Therapeutics, Inc. This relationship is de minimus for Columbia University Medical Center (MH). Drs. Hirano and Carlos Lopez-Gomez are listed as co-inventors of a pending Columbia University patent for gene therapy for TK2 deficiency. The other authors declare no conflicts of interest.

---

**Supplemental figure 1. Weights of female mice.**

Average daily weights in grams for each female treatment group.

**Supplemental figure 2. Rotarod and grip tests in female mice.**

Panel A shows results of rotarod assessments of female mice. Data are expressed as time to fall from the accelerating rotarod. Each symbol represents the average of three tests of a single mouse. For each treatment group, average and standard deviation is represented. Panel B displays bar test (upper limbs) and panel shows grid test (four limbs) in female mice. Data is expressed as strength (measured in g) per weight unit (g). Each symbol represents the average of three tests of a single mouse. For each treatment group, average and standard deviation are represented.

